# Multi-center benchmarking of cervical spinal cord RF coils for 7 T MRI: A traveling spines study

**DOI:** 10.1101/2025.01.13.632825

**Authors:** Eva Alonso-Ortiz, Daniel Papp, Robert L Barry, Kyota Poëti, Alan C Seifert, Kyle M Gilbert, Nibardo Lopez-Rios, Jan Paska, Falk Eippert, Nikolaus Weiskopf, Laura Beghini, Nadine Graedel, Robert Trampel, Martina F Callaghan, Christoph S Aigner, Patrick Freund, Maryam Seif, Aurélien Destruel, Virginie Callot, Johanna Vannesjo, Julien Cohen-Adad

## Abstract

**Purpose:** The depth within the body, small diameter, long length, and varying tissue surrounding the spinal cord impose specific considerations when designing radiofrequency coils. The optimal coil configuration for 7 T cervical spinal cord MRI is unknown and, currently, there are very few coil options. The purpose of this work was (1) to establish a quality control protocol for evaluating 7 T cervical spinal cord coils and (2) to use that protocol to evaluate the performance of 4 different coil designs.

**Methods:** Three healthy volunteers and a custom anthropomorphic phantom (the traveling spines cohort) were scanned at seven 7 T imaging centers using a common protocol and each center’s specific cervical spinal cord coil. Four different coil designs were tested (two in-house, one Rapid Biomedical, and one MRI.TOOLS design).

**Results:** The Rapid Biomedical coil was found to have the highest B_1_^+^ efficiency, whereas one of the in-house designs (NeuroPoly Lab) had the highest SNR and the largest spinal cord coverage. The MRI.TOOLS coil had the most uniform B_1_^+^ profile along the cervical spinal cord; however, it was limited in its ability to provide the requested flip angles (especially for larger individuals). The latter was also the case for the second in-house coil (MSSM).

**Conclusion:** The results of this study serve as a guide for the spinal cord MRI community in selecting the most suitable coil based on specific requirements and offer a standardized protocol for assessing future coils.

## Introduction

While magnetic resonance imaging (MRI) of the human spinal cord (SC) has been a standard clinical procedure for decades, imaging the SC remains challenging due to its small diameter (∼ 1 cm) [1] and depth within the anatomy, which translates into high imaging resolution and signal-to-noise ratio (SNR) requirements. Additionally, its anatomical location near the lungs and major blood vessels makes it highly susceptible to physiological noise, further compromising the achievable SNR. Given that SNR increases almost quadratically [2] with magnetic field strength, the recent emergence of ultra-high field MRI scanners, operating at 7 T and above, has opened up new and exciting opportunities for SC imaging. High-resolution (0.3 x 0.3 mm in-plane) and high SNR [3] SC imaging can facilitate the identification of SC abnormalities [4], detection of SC lesions [5,6] in multiple sclerosis, and visualization and quantification of SC substructures [7].

Notwithstanding its promise, 7 T MRI comes with its own set of challenges, notably the fact that the wavelength of the excitation (B_1_^+^) field is comparable to the size of most anatomical structures of interest. This results in several problems, including short radiofrequency (RF) wave penetration depths and RF wave interference patterns within the body [8]. Third party coil manufacturers and academic research units have rapidly adapted to meet the needs of the 7 T imaging community by developing local transmit (Tx) and receive (Rx) arrays. Options for SC imaging, however, are limited and mostly focused on the cervical levels [9,10]. The small diameter, long length, and varying depth of the SC impose specific considerations when designing RF coils. Moreover, coil designs should be robust to intersubject variability [11], as the depth and curvature of the spine (as well as the surrounding tissues) can vary between individuals. SC RF coil designs must maximize the sensitivity to signals originating from the SC (Rx) and generate B_1_^+^fields (Tx) with sufficient homogeneity along its length to prevent signal bias. The optimal coil configuration needed to achieve this at 7 T is unknown. Design options range from having separate Tx and Rx elements, to sharing the Tx and Rx function for the same array elements (a transceiver array of Tx/Rx elements). The benefit of combining Tx and Rx elements is that of a simplified design, which can also lead to a low number of channels. The latter is of interest considering that most scanners allow for a maximum of 8 Tx channels. Another benefit is that of higher power efficiency, whereas a drawback is the fact that Tx elements in close proximity to the body lead to more steeply varying electric (E) fields, in turn causing a higher specific absorption rate (SAR) [3]. Moreover, the Tx and Rx profiles for loop elements typically have different spatial patterns, making it impossible to optimize the position of elements with respect to the SC for both Tx and Rx performance when using a single element for both functions [3]. Alternatively, if separating Tx and Rx channels, B_1_^+^uniformity can be improved by placing Tx elements further from the anatomy, while sensitivity and the field-of-view (FOV) can be increased by increasing the number and proximity (to the anatomy) of Rx elements. Additional considerations include the choice of rigid vs. flexible coil casings (which can provide increased sensitivity by reducing the distance between the coils and the anatomy) and the geometry of Tx/Rx elements. Proposed geometries include linear arrays of Rx elements [12,13], Tx dipole arrays [14,15], and wrap-around arrays [16], among others (see [9,10] for a review of 7 T SC coil designs).

Given the appropriate hardware interface with the MR system, multi-channel coil arrays can be used in a parallel-transmit (pTx) mode, where each coil element is used to produce an independent and spatially distinct RF field pattern [17,18]. PTx can help address the problem of non-uniform B_1_^+^ due to RF wave interference by optimizing individual channels’ magnitude and phase, such that the overall B_1_^+^field homogeneity is improved over a defined region of interest. The latter is referred to as RF (or B_1_^+^) shimming. For the safe use of RF shimming, virtual observation points (VOPs) [19] are necessary, yet at the time of writing not all coil vendors have made these available. Alternatively, pTx-capable systems can also be used in a virtual single-Tx (1Tx) mode with predefined B_1_^+^phases that are integrated in the hardware or fixed in a coil configuration file (i.e., “CP mode”).

Given the current landscape, researchers seeking to purchase or design a 7 T SC coil are presented with limited options and little guidance for decision making. The purpose of this work was twofold: (1) To establish a quality control (QC) protocol for evaluating 7 T SC coils’ Tx (B_1_^+^) and Rx (SNR, g-factor) characteristics; and (2) To use that QC protocol to evaluate the performance of 4 different cervical SC coil designs. The included coils were situated at 7 different MR centers and evaluated in the same group of 3 volunteers and an anthropomorphic phantom that traveled to each of the participating centers (the traveling spines cohort). The results obtained in this study could serve as a guide for the SC MRI community to choose the most suitable coil based on their requirements.

## Material and methods

### Participating Centers and RF coils

The participating MRI centers were: Centre de Résonance Magnétique Biologique et Médicale (CRMBM, Marseille), University College London (UCL, London), Montreal Neurological Institute (MNI, Montreal), Athinoula A. Martinos Center for Biomedical Imaging (MGH, Boston), Max-Planck Institute for Human Cognitive and Brain Sciences (MPI, Leipzig), Norwegian University of Science and Technology (NTNU, Trondheim), and Mount Sinai Biomedical Engineering and Imaging Institute (MSSM, New York).

The comparison was based on 4 different 7 T SC MRI coil designs of different surface array geometries, covering different FOVs. The coils include 2 in-house built coils (NeuroPoly Lab and MSSM) and 2 commercial coils (Rapid Biomedical and MRI.TOOLS). Three of the coils have pTx (and therefore RF-shimming) capabilities (NeuroPoly, Rapid Biomedical, and MRI.TOOLS), 3 are designed for cervical spine imaging, and 1 is designed for posterior fossa and cervical spine imaging (MSSM). Figure 1A provides an overview of the included sites and the coil element layouts for the employed coils.

**Figure 1:**
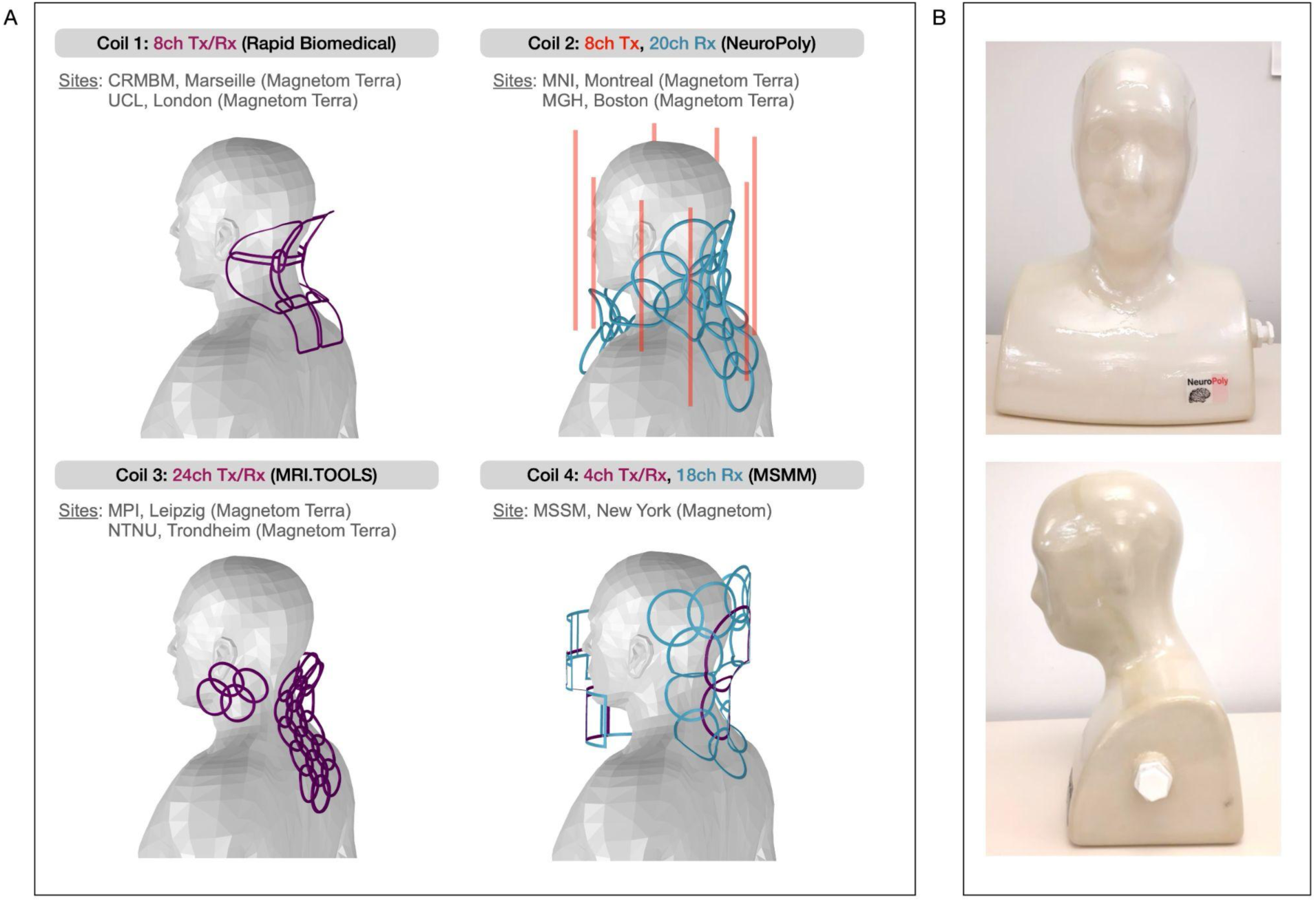
(A) Coil element layout for the 4 tested designs: Coil 1 (Rapid Biomedical), Coil 2: NeuroPoly (in-house), Coil 3: MRI.TOOLS, Coil 4: MSSM (in-house). Tx-only coils are shown in red, Rx-only in blue, and Tx/Rx in purple. The site(s) possessing a coil design that was included in this comparative study is indicated below the coil description, as well as the corresponding 7 T system. (B) Spinoza V6 phantom

#### Coil 1 (Rapid Biomedical [20])

This coil includes a transceiver array of 8 elements spread on a posterior surface that cradles the subject’s neck. The central element is centered at cervical vertebrae 3-4 (C3-C4) and the coverage in the head to foot (H/F) direction extends roughly from the cerebellum to the thoracic vertebra 1 (T1). By default, a power splitter combines all of the elements for transmission (1Tx), with an identical RF waveform delivered to each element (used in this study). With the availability of a virtual observation points (VOPs) file, and by removing the power splitter, all 8 channels can be used for pTx transmission [21].

#### Coil 2 (in-house from NeuroPoly [22])

The coil contains 8 dipole elements for transmission, one anterior, three posterior, and four lateral, with extensive coverage along the H/F axis. The coil includes 20 circular Rx loops, divided into two groups: an array of 15 loops on the posterior and lateral parts of the casing and 5 loops which are integrated into a detachable and functionally independent assembly that is placed anterior to the neck and the upper chest. The latter 5 loops were intended to improve the overall sensitivity of the coil, especially in the lower part of the cervical SC, where the distribution of the posterior 15 Rx loops is less dense. The distance between the anterior, detachable casing and the subject can be adjusted, allowing one to better adapt the coil to a large variety of subject sizes, thereby improving the SNR. The coil used at the MGH site is a reproduction of the one used at the MNI. They have the same Tx and Rx elements, with the same geometry and electrical design. However, the overall mechanical design is slightly different, leading to the Tx coil wiring and its dielectric environment (supports) to change as well. These changes result in the S-parameters being slightly different from those of the MNI coil. This coil was used in CP-mode.

#### Coil 3 (MRI.TOOLS)

This 24 channel coil is designed for cervical SC imaging and comprises: (1) A 16-channel phased array with three rows, designed to contour the curvature of the neck. This array extends from the back of the head to approximately T3/T4. (2) Two 4-channel phased arrays, positioned on either side of the head. The coil’s 24 channels are configured for Tx/Rx functionality and grouped into six sets of four channels. These groups are managed via six 4-way power splitters. For flexibility, each splitter can operate independently with the pTx array. Alternatively, all splitters can be combined into a single transmit channel using an additional 6-way power splitter (used in this study).

#### Coil 4 (in-house from MSSM [23])

This coil has a 2 panel design, designed to boost SNR in the center of the subject’s neck with cervical SC and brainstem coverage. The coil consists of 4 Tx/Rx channels and 18 Rx channels that fully encircle the head and neck. Two rectangular Tx/Rx elements are placed anterior and two circular elements are placed posterior to the head and neck. The anterior panel contains six additional rectangular Rx elements and the posterior panel contains 12 circular Rx elements. While this coil is not pTx capable in its current implementation, it would be possible to convert it to pTx by bypassing the power dividers and replacing plugs. The coil does however have a rudimentary phase-only adjustment, achieved by varying the length of the coaxial patch cable between the anterior and posterior halves.

### Subject Measurements

Three healthy volunteers (Subject 1 [37 years, 195 cm, 110 kg, M], Subject 2 [26 years, 155 cm, 40 kg, F] and Subject 3 [45 years, 170 cm, 77 kg, M]) and a custom anthropomorphic phantom (Figure 1B) traveled to each participating center and were scanned using the center’s corresponding coil. Informed consent was given prior to each in-vivo scanning session in accordance with each center’s ethics committee.

### Phantom Design

We built a phantom (“Spinoza V6”) [24] for this study (Figure 1B) with a shape based on a 3D average head model derived from MPRAGE scans obtained from 22 adult males [22] that was 3D printed using PLA 2.0 EcoTough™. The phantom was filled with a 15.4 L solution containing demineralized water (641 mL/L), sugar (513 g/L), salt (16.5 g/L), mouthwash (40 ml/L), agar (7.5 g/L). This formula was based on a recipe from Dual et al. [15], with the addition of mouthwash (which has 4% Listerine, an antibacterial agent). A conductivity of 0.74 S/m and a permittivity of 65.8 were measured at 300 MHz with an Agilent 85070E dielectric probe. These parameters mimic the average coil loading of the biological tissues that make up our region of interest. Spinoza V6 was used to acquire phantom scans at all participating sites. More information on the phantom can be found here: https://github.com/neuropoly/phantom-spinoza

### Protocol Design and Coil Evaluation Criteria

Protocol parameters were harmonized as much as possible across Siemens Magnetom 7 T and Magnetom Terra systems for all sequences. The protocol PDF printouts and executable (.exar1) files used at each site for the phantom and each volunteer can be found here: https://github.com/spinal-cord-7t/coil-qa-protocol. A detailed QC standard-operating-procedure (SOP) for phantom and in-vivo scans, that was followed by all scan operators can also be found at that same url. Here we describe the sequences that were included in the study, the rationale for using specific sequences and acquisition parameters, particularities of the acquisition for said sequences, and related post-processing. The SOP provides step-by-step instructions for acquiring data using both 1Tx/CP-mode and RF-shimming. However, all of the below-described scans were acquired without RF-shimming.

#### Anatomical scanning

##### Sequence

MP2RAGE [25]

##### Rationale

MP2RAGE scans were included in the in-vivo protocol for the purpose of segmenting the SC and comparing high quality SC images across coils.

##### Acquisition (based on [22])

3D coronal acquisition with 0.7 mm isotropic resolution, TR = 5000 ms, TE = 2.15 ms, TI1/TI2 = 700/2400 ms, flip angle (FA)1/FA2 = 4^0^/5^0^, FOV = 260 x 171 mm^2^, 192 slices, total acquisition time (TA) = 8m47s.

#### B_1_^+^ mapping

##### Sequences

2D DREAM [26] and 2D presaturated turboFLASH [27]

##### Rationale

Two B_1_^+^mapping sequences (DREAM [28] and presaturated turboFLASH (TFL) [29]) were included in the protocol to balance the strengths and weaknesses of each sequence.

For both B_1_^+^mapping sequences, the FOV was set to the maximum size allowed by the scanner to evaluate B_1_^+^profiles across the entirety of the coils’ coverage, as well as to compare the extent of each coil’s coverage. Sagittal acquisitions were included in order to maximize H/F coverage while minimizing acquisition time and potential motion artifacts (which are less common in the left(L)/right(R) direction). The phase encoding direction was set to anterior(A)/posterior(P). For the DREAM sequence, the slice thickness (2.5 mm) and in-plane resolution (4.1 x 4.1 mm^2^ and 2.5 x 2.5 mm^2^) were selected in order to provide adequate SNR while still minimizing partial volume effects with cerebrospinal fluid (CSF). Due to the inherent higher-SNR of the TFL images, compared to those obtained with the DREAM sequence (due to the use of stimulated echoes), a higher in-plane resolution was used. Additional DREAM scans were acquired with varying reference voltages to use the approach proposed in [30] and a smaller FOV size and a higher image resolution in order to mitigate relaxation throughout the readout train.

##### Acquisition

First, the B_1_^+^field was mapped with a TFL sequence using the reference voltage obtained with the scanner’s built-in voltage optimization method. The latter is designed to find the voltage required to obtain a 180° flip angle distribution using a 1 ms hard pulse, measured over 3 axial slices (at z = -5, 0, +5 cm) across the entire FOV. The reference voltage required to reach the desired flip angle was calibrated by manually choosing an ROI on the TFL FA map within a region approximately corresponding to the cervical level for which the coil was optimized (Rapid Biomedical: this coil is optimized for the cervical SC, however C3 was used as a landmark (approximately the center of the coil); NeuroPoly in-house: C6/7; MRI.TOOLS: this coil is optimized for C1-T1, however, C5/6 and C2/3 were used as landmarks at MPI and NTNU, respectively; MSSM in-house: C3/4) or the center of the FOV at the midline (phantom scans). This reference voltage, termed “optimal reference voltage, *V_opt_*” was used for all subsequent scans, with the exception of DREAM sequences deliberately acquired at different reference voltages.

For some coil and volunteer combinations, the reference voltage required to attain the prescribed saturation FA when using presaturated TFL at the prescribed region either exceeded the coil’s hardware limit or led to clipping of RF pulses. In those instances, the reference voltage was reduced to avoid RF pulse clipping (all three subjects when using the MRI.TOOLS coil at MPI) or set to the maximum permissible (Subjects 1 and 3 when using the MRI.TOOLS coil at NTNU, and Subject 1 when using the MSSM coil after having reduced the saturation RF pulse FA for TFL to avoid pulse clipping).

The B_0_ adjustment volume was set to be the same as that of the FOV and 3D distortion correction was used.

TFL and DREAM acquisition parameters are detailed in Supplementary Table 1.

#### B_1_^-^ mapping

##### Sequences

The coilQA package (C2P available for Magnetom centers running VB and VE platforms: https://www.nmr.mgh.harvard.edu/c2p, available as a product sequence for Terra centers) was selected for assessing coil Rx performance. The image reconstruction implementation of this 2D sequence conveniently provides SNR, g-factor and noise statistic maps. Lastly, a single 3D gradient-echo (GRE) acquisition was included in the protocol so that individual channels could be qualitatively assessed (individual coil images were not combined).

##### Rationale

coilQA images were acquired sagittally with as large a FOV as possible in the H/F direction to assess the full extent of each coil and with a smaller FOV that is more representative of actual acquisitions. Sagittal scans were acquired with the phase encode direction set to H/F in order to avoid motion-related artifacts. The image resolution was carefully selected to maximize SNR, while also minimizing susceptibility-induced signal loss around the intervertebral junctions. An additional, high-resolution, axial coilQA scan was acquired to reflect realistic scan scenarios that may be used for quantitative MRI, such as spinal cord area measurements.

##### Acquisition

To minimize susceptibility-induced signal loss within the spinal cord, the size of the B_0_ shim adjustment volume along the A/P and L/R directions was adjusted to encompass the spinal cord and as few surrounding tissues as possible for all coilQA acquisitions. coilQA acquisition parameters are detailed in Supplementary Table 2.

### Processing

DREAM FA maps acquired at *V_opt_*, (2/3)*V_opt_*, and 1.5*V_opt_* were thresholded to exclude voxels with FA less than 20° or greater than 50°. Each one of those FA maps were converted to B_1_^+^maps in units of nT/V. The three B_1_^+^maps were finally averaged to result in one DREAM B_1_^+^map. The TFL FA maps were also converted to B_1_^+^in units of nT/V.

#### Phantom

To quantify B_1_^+^and SNR, an approximate SC mask was generated by first selecting a representative human anatomical scan and overlaying this scan onto the CRMBM phantom TFL scan. Pointwise labels were then created along the SC of the human scan and interpolated onto the phantom image. Lastly, a cylindrical mask of 10 mm diameter was generated along the centerline joining the pointwise labels. See Figure 2 for a visual summary of these steps.

**Figure 2:**
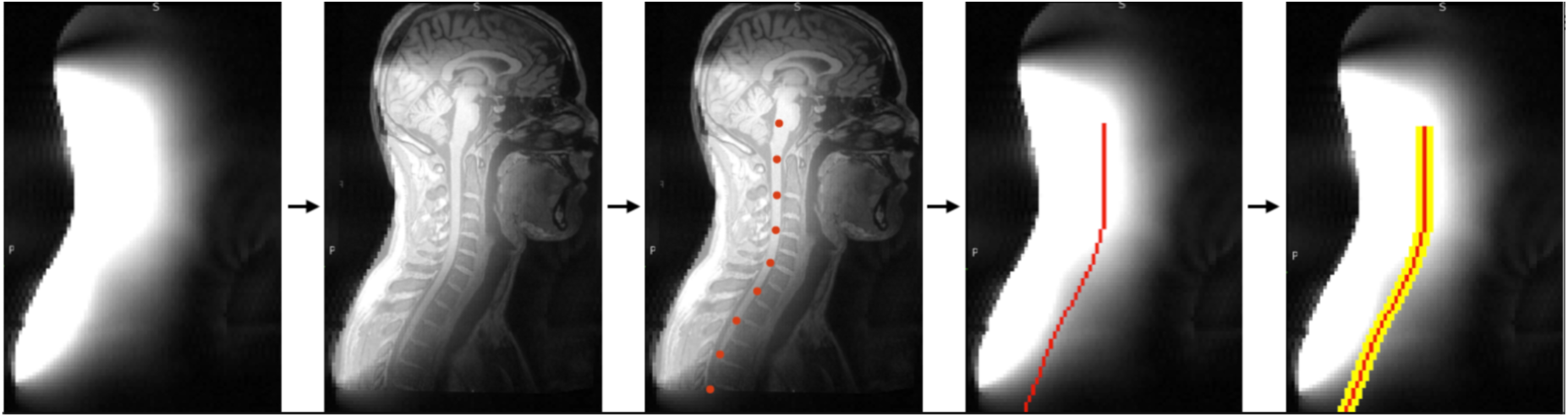
Phantom SC mask creation workflow. The leftmost image is the CRMBM phantom scan that was coregistered onto a representative human anatomical scan. The red dots represent SC labels that were joined (red ROI) to form a path that was dilated, resulting in a phantom SC mask (yellow ROI), covering roughly from brainstem to T5-T6 vertebral level.

B_1_^+^and SNR data were co-registered across sites, such that the same (above-described) SC mask could be used for all sites. Co-registration to a reference site (CRMBM) was achieved using labels that were created for each site at the posterior tip and inflection point of the phantom (see Figure 3).

**Figure 3:**
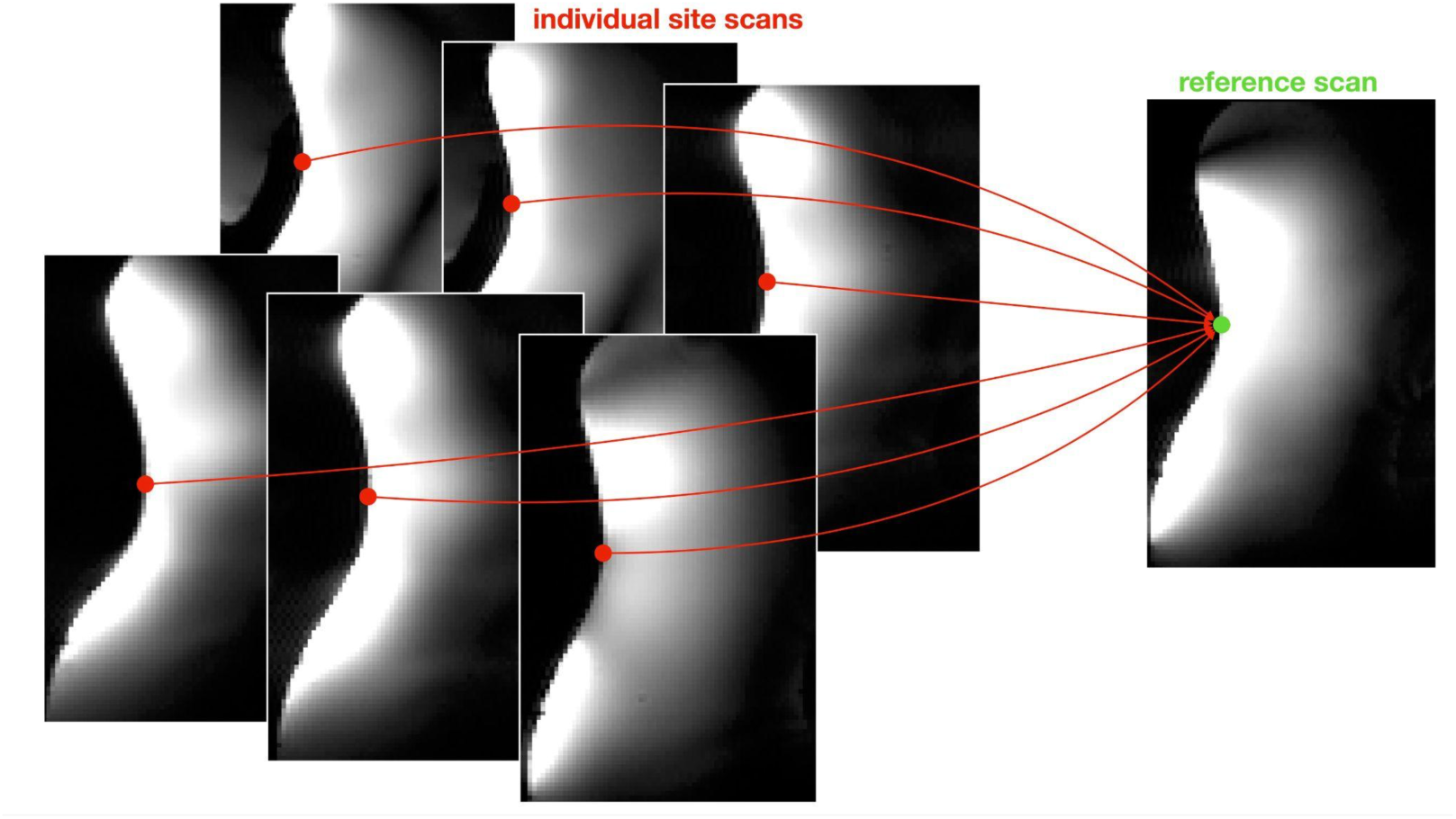
Co-registration of phantom scans across sites

B_1_^+^and SNR were each averaged in the axial plane within the SC mask, and those averaged SNR values were corrected for B_1_^+^field effects by dividing by sin(FA_GRE,meas_) [31], where FA_GRE,meas_ was obtained from the TFL B_1_^+^scan as follows: FA_GRE,meas_ = FA_GRE,requested·_(FA_TFL,meas_/FA_TFL,requested_). By normalizing, the SNR value achieved with FA = 90° was extrapolated (SNR_90_).

#### In-Vivo

The Spinal Cord Toolbox (SCT) [32] was used to automatically segment the SC and label vertebral disks on MP2RAGE UNI images. SC masks and labeled vertebral disks were manually corrected where needed.

The SC was segmented using SCT on the magnitude images of the TFL and DREAM scans and the large sagittal-FOV SNR images. Those SC segmentation masks were used to facilitate the registration of TFL and DREAM FA and SNR maps onto MP2RAGE UNI images. The inverse co-registration transform was applied to MP2RAGE UNI-derived SC masks and labels to bring them into the space of the TFL and DREAM FA and SNR maps. The vertebral-level (non-zero) average TFL and DREAM B_1_^+^and SNR measurements within the SC mask were then extracted between C1 and T2 (roughly corresponding to the greatest coverage available among the included coils) and smoothed using SciPy’s “uniform_filter1d” filter. The latter SNR measurements were subsequently corrected for B_1_^+^field effects using the approach described above for phantom data.

## Results

### B_1_^+^ mapping

TFL B_1_^+^maps of Spinoza (Figure 4A) and human subjects (Figure 4B) showed a diverse range of B_1_^+^efficiency profiles. As shown in the bottom left of Figure 4A, the CRMBM/UCL (Rapid Biomedical) coil shows the highest peak B_1_^+^efficiency (i.e., the ratio of produced B_1_^+^amplitude per transmit voltage), whereas the MNI/MGH (NeuroPoly) coil had the greatest coverage along the H/F + A/P directions for phantom acquisitions. The SC-averaged TFL B_1_^+^from C1 to T2 for each subject at each site are displayed in Figure 6. Here the trends observed in Spinoza are mostly reflected, with the CRMBM/UCL coil showing the highest peak B_1_^+^efficiency. The subject-averaged (solid black line in Figure 6A) TFL B_1_^+^along the SC was averaged across vertebral levels for each site and found to be: 40±9 (CRMBM), 37±7 (UCL), 24±3 (MNI), 22±3 (MGH), 18±2 (MPI), 21±1 (NTNU), 28±5 (MSSM) nT/V. When considering DREAM-measured B_1_^+^efficiency (Figure 5 and Figure 6B), the maps showed some visible differences (compared to TFL), while trends along the SC were somewhat consistent with TFL. The subject-averaged DREAM B_1_^+^along the SC, averaged across vertebral levels for each site, was: 39±6 (CRMBM), 35±4 (UCL), 32±3 (MNI), 22±2 (MGH), 26.3±0.3 (MPI), 27±2 (NTNU), 30±3 (MSSM) nT/V. The root-mean squared error (RMSE) between TFL and DREAM estimates across coils was 5 nT/V.

**Figure 4:**
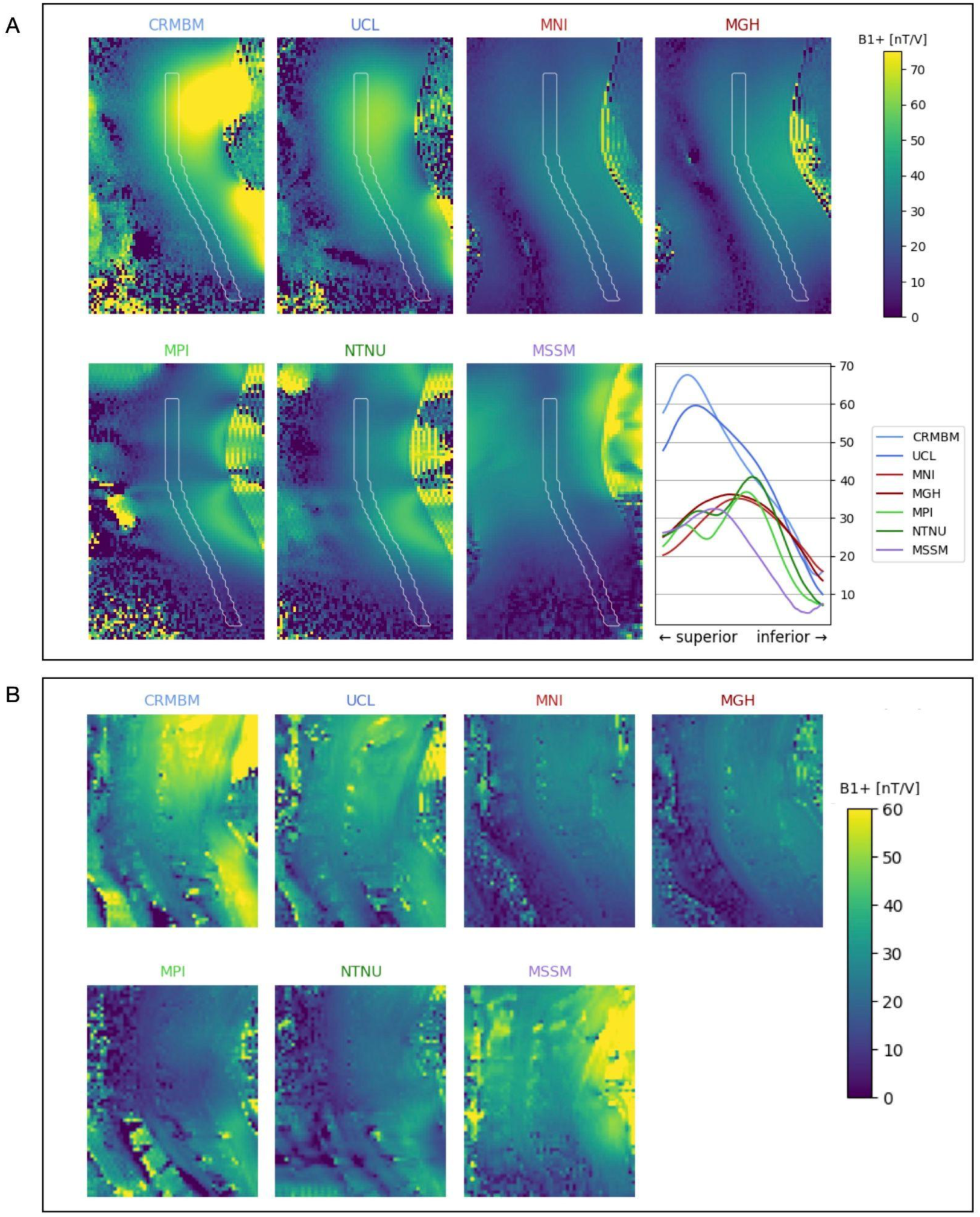
(A) Spinoza TFL *B*_1_^+^maps at each participating site. Last figure: measured *B*_1_^+^along the SC mask (white outline) for each site. (B) In-vivo TFL *B*_1_^+^maps at each participating site for the same representative subject (Subject 3).

**Figure 5:**
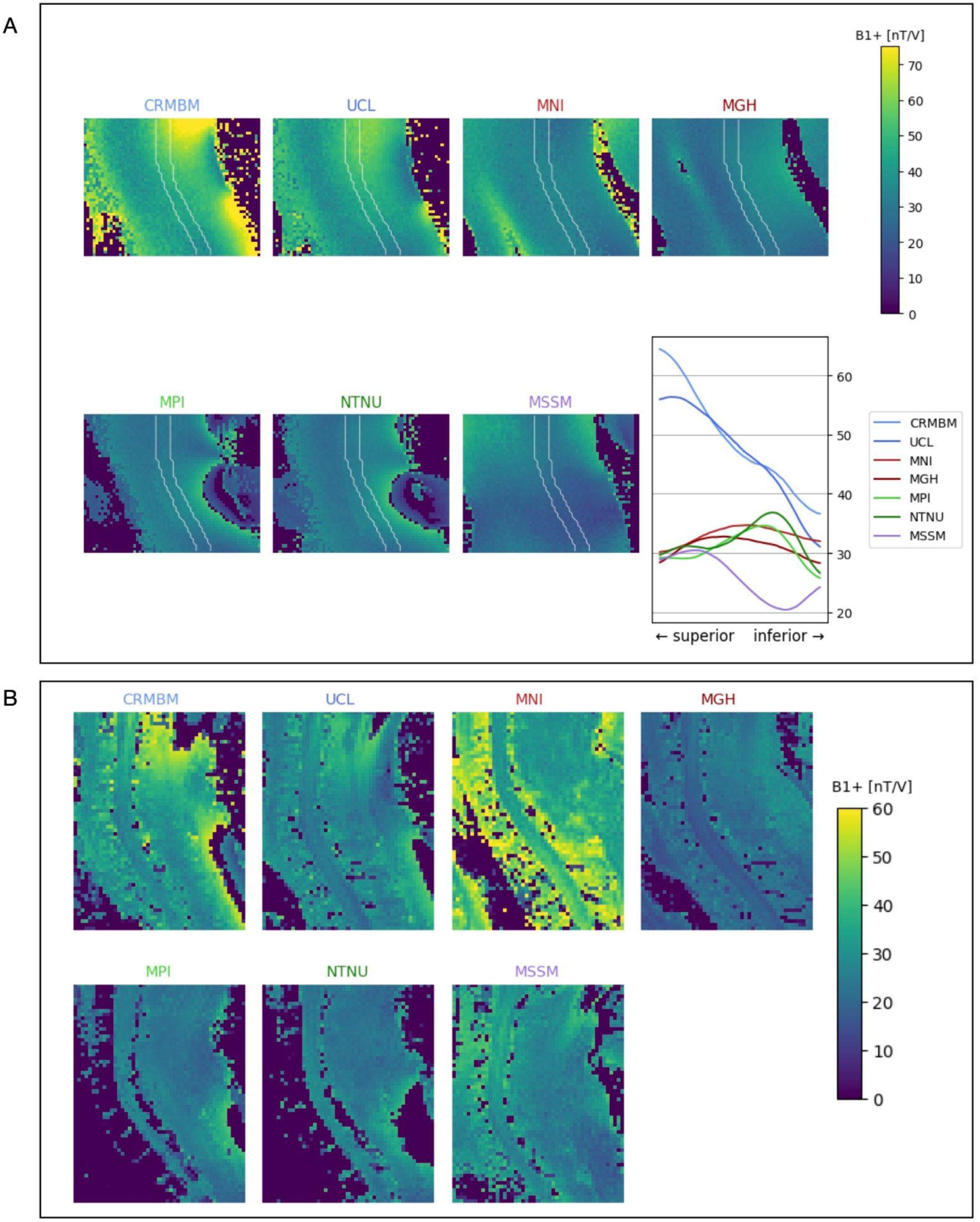
(A) Spinoza DREAM *B*_1_^+^maps at each participating site. Last figure: measured *B*_1_^+^along the SC mask (white outline) for each site. (B) In-vivo DREAM *B*_1_^+^maps at each participating site for the same representative subject (Subject 3).

**Figure 6:**
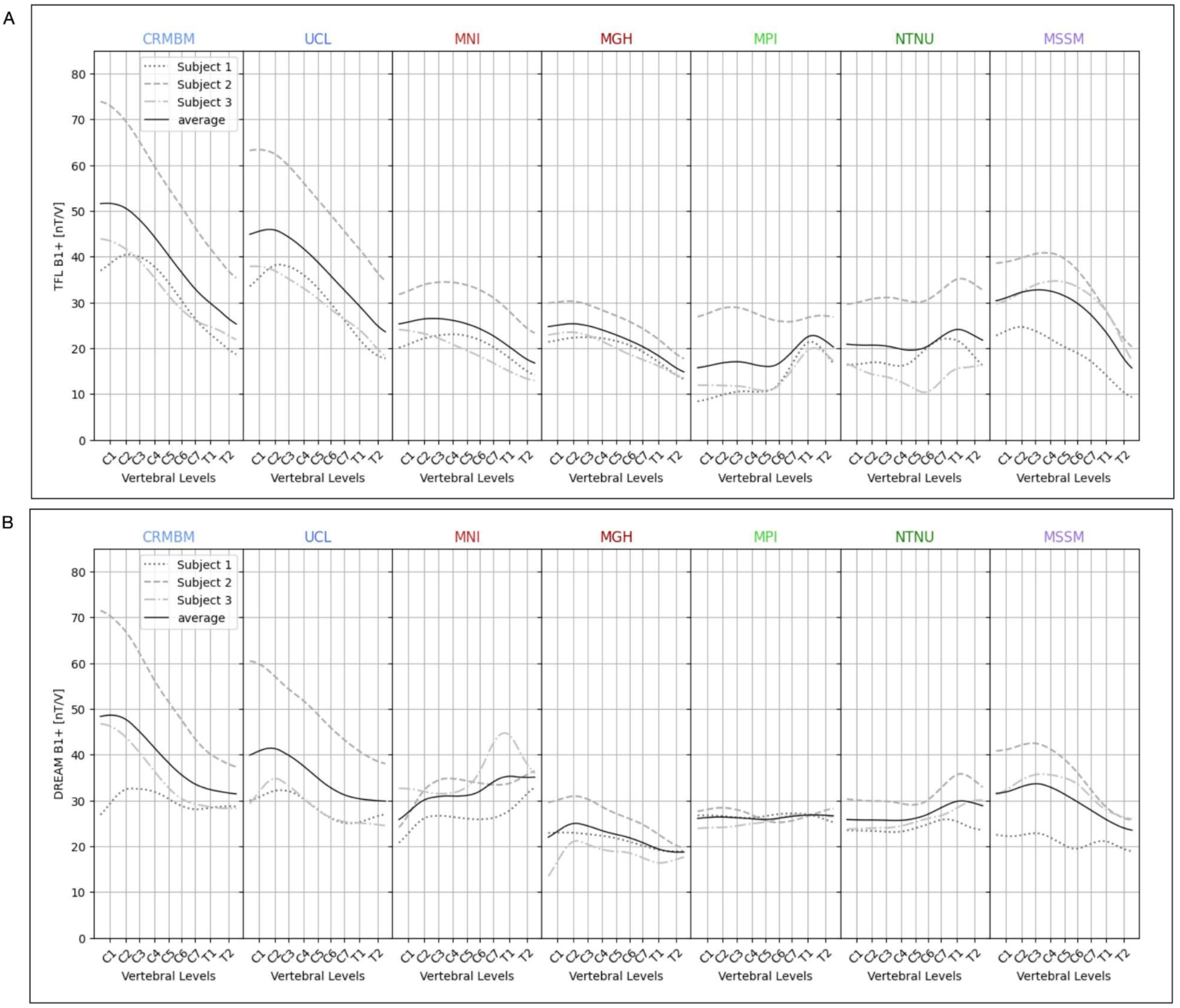
(A) TFL *B*_1_^+^along the SC from C1-T2 across sites for each subject. (B) DREAM *B*_1_^+^along the SC from C1-T2 across sites for each subject.

Generally TFL and DREAM B1+ were in agreement both in terms of measured values, and trends along the vertebral levels. One exception was the MNI coil; DREAM B1+ along the cord for that coil displayed a reversed trend, compared to TFL measurements (with high to low B_1_^+^along the H/F direction for the latter).

One can also appreciate that the smallest subject (Subject 2) presented the highest B_1_^+^efficiency values along the SC for all coils (barring DREAM MNI data). To quantify the consistency across subjects, the standard deviation (SD) across subjects of the SC-averaged B_1_^+^was computed for TFL/DREAM: 10.9/9.9 (CRMBM), 10.1/9.6 (UCL), 5.4/3.7 (MNI), 2.9/3.4 (MGH), 6.4/0.7 (MPI), 7.5/3.1 (NTNU), 6.6/6.2 (MSSM). The discrepancy between measured B_1_^+^in the smaller female subject and the two male subjects was greatest for the CRMBM coil (SD_TFL_ = 10.9 and SD_DREAM_ = 9.9). When considering both TFL and DREAM data, the MGH coil appeared to have the smallest discrepancy (SD_TFL_ = 2.9 and SD_DREAM_ = 3.4). If considering only the DREAM B_1_^+^maps, the MPI coil appeared to lead to comparable values for larger/smaller subjects (SD_DREAM_ = 0.7).

Regarding the variability of B_1_^+^along the SC (Figure 7A), the NTNU coil displayed the lowest subject-averaged coefficient of variation (CoV) and the CRMBM coil the highest. It is noteworthy that the TFL B_1_^+^CoV data for MPI and NTNU appear to differ more than their coil-pair counterparts (CRMBM/UCL and MNI/MGH). This may be related to the fact that at the MPI site data for Subjects 1, 2 and 3 were not acquired using *V_opt_* to avoid RF clipping (Subject 1: 500 V used instead of 2079 V, Subject 2: 510 V used instead of 545 V, Subject 3: 510 V used instead of 1013.5 V). On the other hand, at the NTNU site, Subjects 1 and 3 were also not acquired using, but rather at a reference voltage respecting the coil’s hardware limit (Subject 1: 550 V used instead of 1003 V, Subject 3: 550 V used instead of 1037 V).

**Figure 7:**
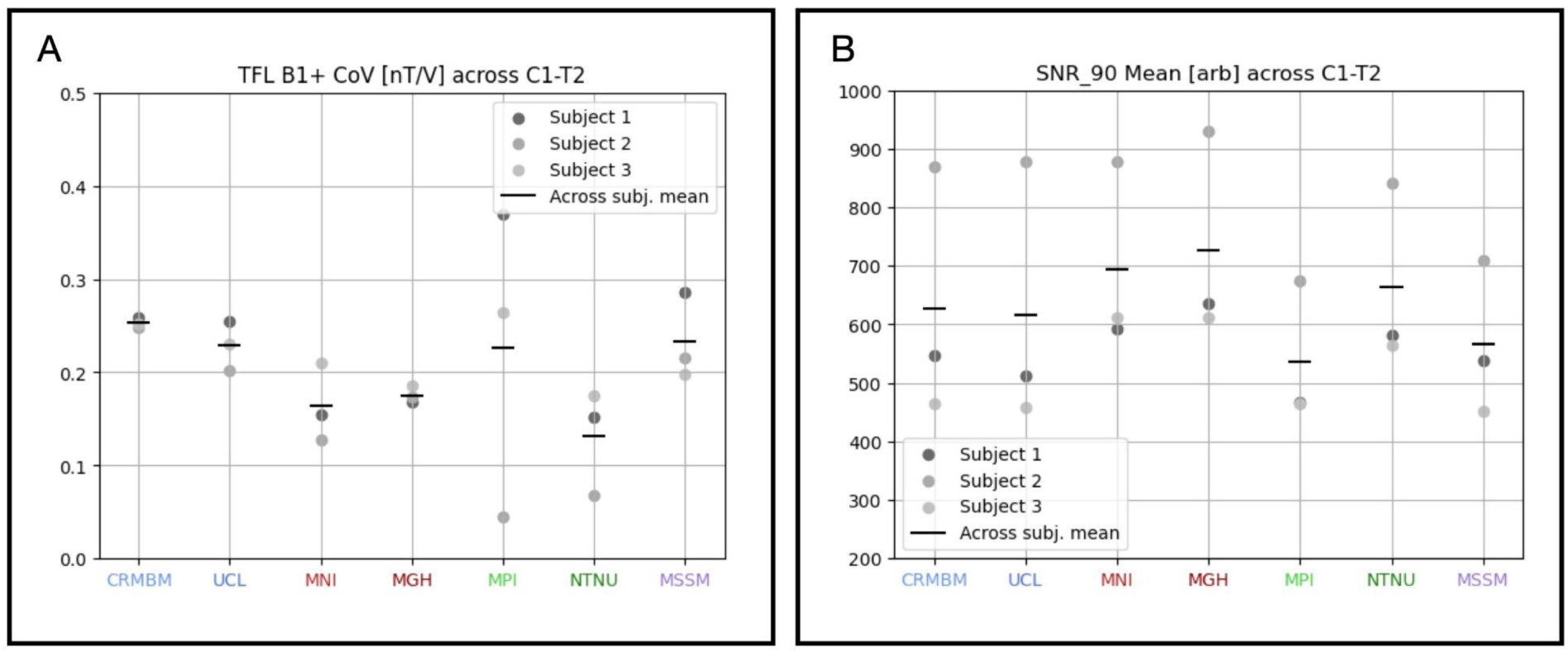
(A) Coefficient of variation (CoV) across C1-T2 in the spinal cord of TFL B_1_^+^for each subject, across sites. (B) SNR_90_ averaged across vertebral levels in the spinal cord. The average B_1_^+^CoV and SNR_1_^+^across subjects is shown as a black horizontal line.

### B_1_^-^ mapping

SNR maps in the Spinoza phantom (Figure 8A) revealed varying degrees of sensitivity, with the MGH coil having the highest peak SNR_90_ along the SC phantom mask (last subplot in panel A). The MGH/MNI coil led to the greatest coverage in the H/F + A/P directions. Figure 8B shows SNR maps of one representative subject at each site and Figure 9 shows SNR_90_ from C1 to T2, for each subject at each site. The subject-averaged (solid black line in Figure 9) SNR_90_ along the SC was averaged across vertebral levels for each site (Figure 7B) and found to be: 627±40 (CRMBM), 617±25 (UCL), 694±63 (MNI), 726±40 (MGH), 535±112 (MPI), 663±59 (NTNU), 567±71 (MSSM) nT/V.

**Figure 8:**
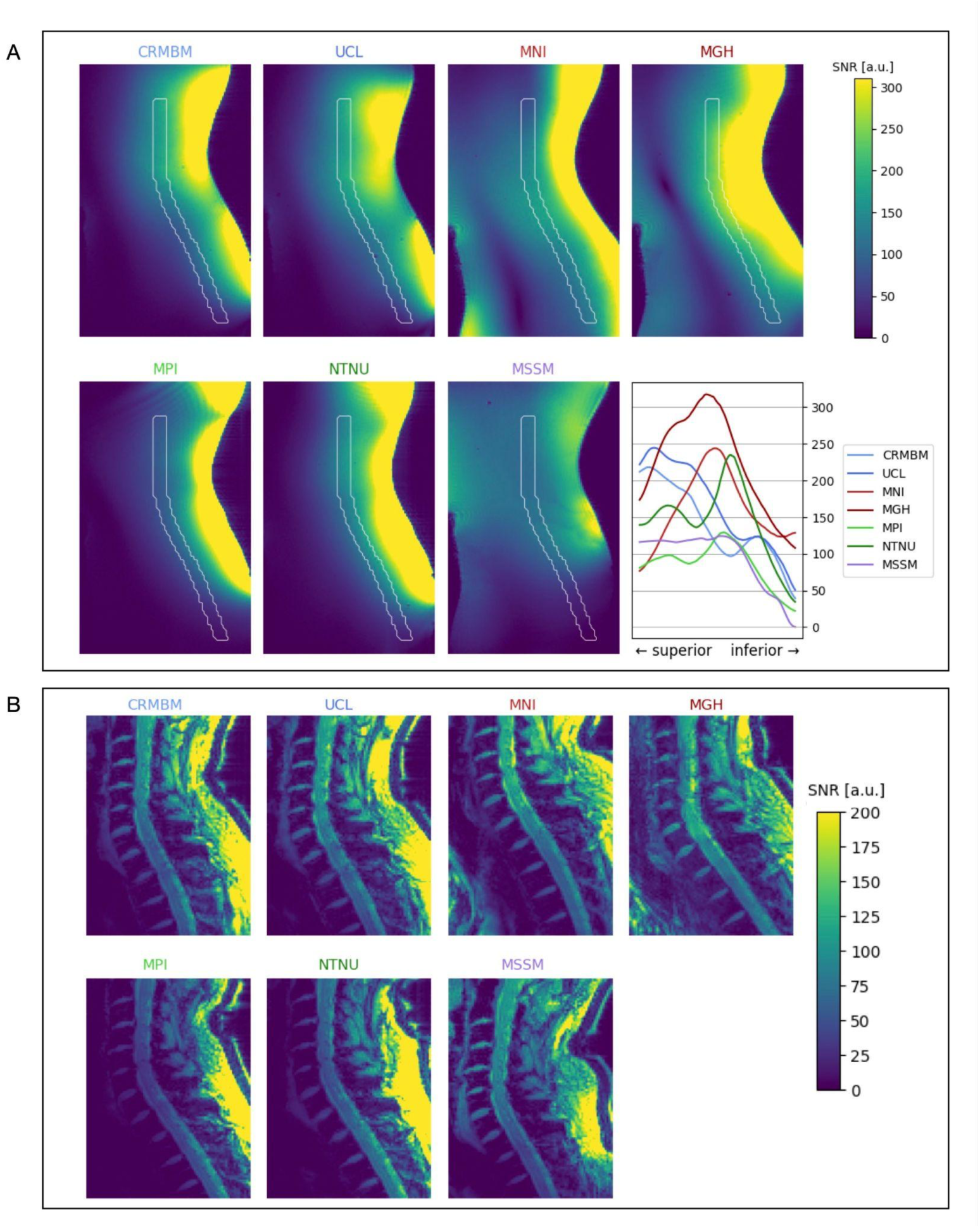
(A) SNR maps measured with the Spinoza phantom at each participating site. The last figure includes the measured SNR_90_ along the SC mask (white outline in the B1+ maps) for each site. (B) SNR for a single representative subject across all sites (Subject 3).

**Figure 9:**
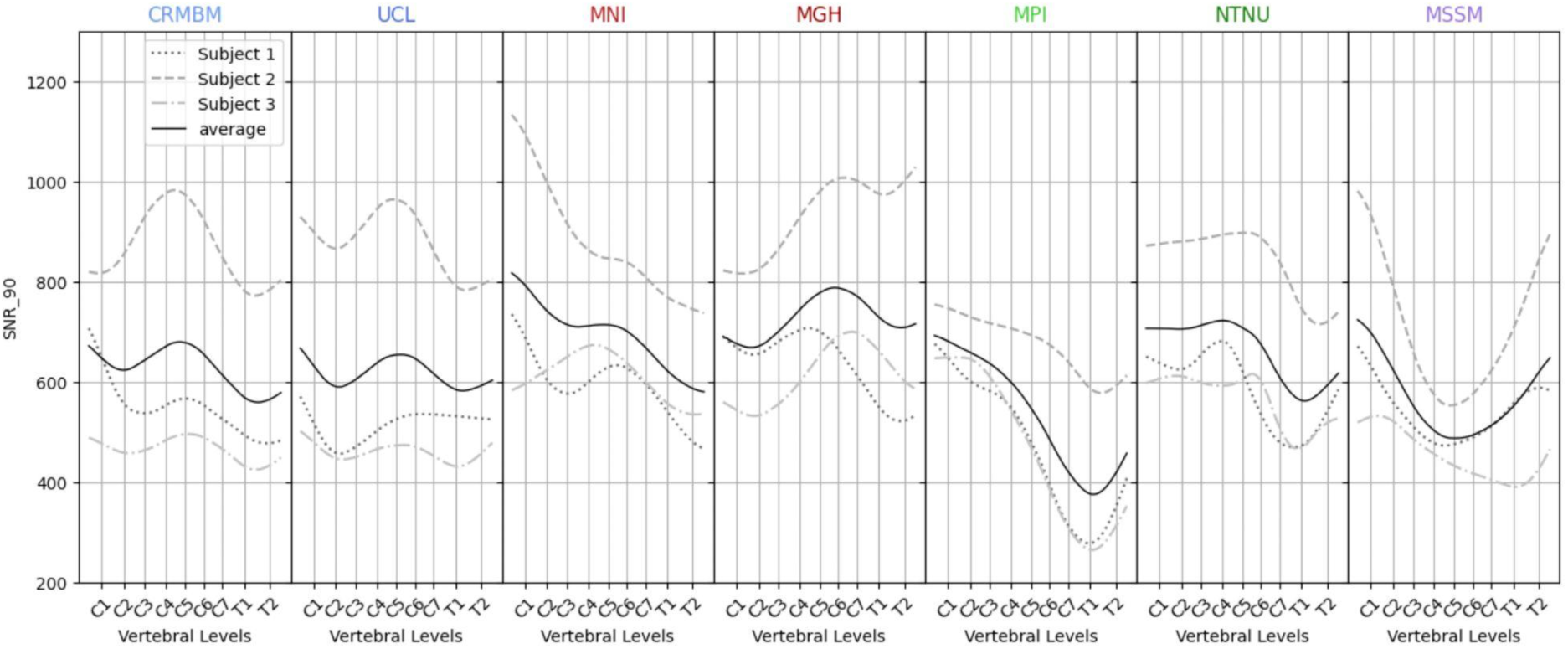
SNR_90_ along the spinal cord from cervical level 1 (C1) to thoracic level 2 (T2) across sites for each subject.

Phantom 1/g-factor maps corresponding to sagittal small-FOV coilQA acquisitions with SENSE undersampling acceleration rates R_H/F_ x R_A/P_ = 2 x 2 are shown in Figure 10A (except for the MSSM site for which there were some unresolved data corruption issues). Average 1/g-factors were measured in a rectangular ROI in the center of the image, corresponding to the central slice. The highest measured 1/g-factor (and, hence, lowest noise amplification) corresponded to the MNI coil (average of 0.967) and the lowest corresponded to the UCL coil (average of 0.851). The percent difference in the average measured 1/g-factor between the MNI and UCL coils was 13.6% for R_H/F_ x R_A/P_ = 2 x 2. In Figure 10B, results are shown for a single representative subject across all sites. Similar to phantom images, the greatest noise amplification was observed for the CRMBM/UCL coil (likely due to the lack of anterior Rx channels). Phantom and in-vivo 1/g-factor maps corresponding to axial coilQA acquisitions with SENSE [33] undersampling acceleration rates R_L/R_ = 2 are shown in Supplementary Figure 1. Here, the CRMBM/UCL coil again displayed the greatest noise amplification (1/g-factor=0.78), whereas the MPI/NTNU coil displayed the lowest (1/g-factor=0.93). The percent difference in the average measured 1/g-factor between the MPI and CRMBM coils was 19.2% for R_L/R_ = 2 in the phantom.

**Figure 10:**
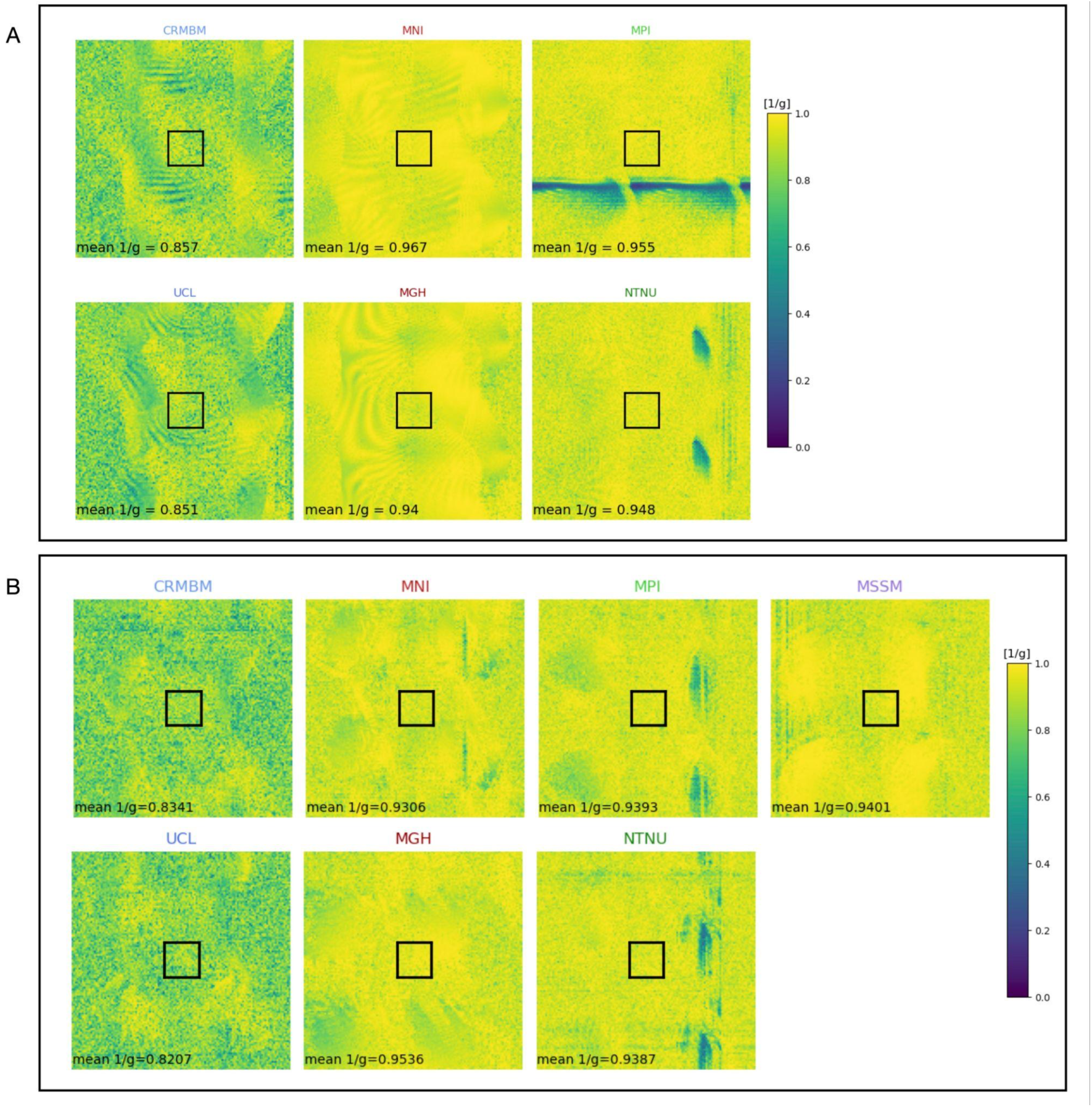
(A) 1/g sagittal maps with R_F/H_ x R_A/P_ = 2 x 2 of Spinoza at each site, except for MSSM. (B) 1/g factor maps with R_F/H_ x R_A/P_ = 2 x 2 for a single representative subject across all sites (Subject 2).

### Additional evaluation criteria

Most coils were rated acceptable by volunteers in terms of patient comfort, with the following exceptions (which were reported by all three volunteers):

- When scan times exceeded 60 minutes, volunteers reported discomfort in the back of the head in the NeuroPoly coil, due to the shape of the coil.
- The Rapid Biomedical coil lacks head support, impeding immobilization and padding. Volunteers reported having to hold their heads straight and still, as there was no support on either side to allow for relaxed positioning.

## Discussion and Conclusions

This study presents a comprehensive QC protocol for evaluating 7 T spinal cord MRI coils, and an assessment of 4 different SC coil designs. We believe that our QC protocol for 7 T SC MRI coils can help the community expedite the coil development process. By adopting a unified consensus on benchmarking criteria, comparisons among various coil designs can be streamlined. This standardization can not only accelerate progress, but also facilitate more efficient assessment of 7 T SC image quality.

Regarding transmit performance, B_1_^+^efficiency, uniformity, and coverage were considered, while SNR and g-factor measurements were used as receive metrics. The proposed QC protocol covers additional aspects of coil performance that have not been presented in the body of this manuscript, but can either be found in the Supplemental Materials (individual channel GRE images for all coils, Supplementary Figures 2-8) or on the following data repositories: https://openneuro.org/datasets/ds005025 (human dataset) and https://openneuro.org/datasets/ds005090 (phantom dataset). The QC protocol SOP also covers the acquisition of QC data with RF-shimming (using the Shimming-Toolbox [34]). Due to time restraints and technical challenges, we were unable to acquire QC data with RF-shimming in this study.

With regards to data processing, the curation of our Brain Imaging Data Structure (BIDS) dataset was not straightforward. We worked with several niche sequences, and some are not yet fully covered by BIDS. The recent qMRI BIDS extension was helpful [35], but there is still work to do to ensure that less common use cases are considered in BIDS. The availability of research sequences across scanner models and software versions presented some difficulties. Not all sequences resulted in equal outputs; for example, the TFL B_1_^+^mapping sequence outputs an anatomical scan on Magnetom Terra systems running VE12U, but not on Magnetom systems running VB17. This anatomical scan was used for segmentation purposes, and its omission presented data processing challenges (in this case, manual segmentations were required). Another challenge was the fact that SCT’s segmentation methods have not been designed for 7 T data, and can be particularly difficult to use with uncommon contrasts (MP2RAGE UNI images, B_1_^+^maps, etc.). Nevertheless, SCT’s newest contrast-agnostic method (sct_deepseg) performed reasonably; only 7 (out of 63) of the datasets required manual corrections. Vertebral labeling was also not reliable on MP2RAGE data and had to be done manually. Recent work [36] should overcome that issue. Finally, it should be noted that the SC is a small (∼1 cm in diameter) structure, hence extracted metrics are prone to strong partial volume effects. This was mitigated by averaging across vertebral levels, smoothing curves, and using soft (non-binary) segmentations [37].

Some of the limitations of this study include the fact that no quantitative comparisons of SAR were made, although this is an important criteria when using sequences with adiabatic inversion pulses. Furthermore, RF-shimming performance was not assessed for pTx-capable coils. Also, it should be noted that this study was limited to 4 different coil designs. Commercial cervical SC coil designs for 7 T are also available from Nova Medical [38] and Quality Electrodynamics [39]. Various additional in-house designs [3,13,15,40–43] have also been proposed, which could not be included in our study.

When it comes to interpreting B_1_^+^mapping results, one must bear in mind that both TFL and DREAM sequences included in this study each have their own strengths and weaknesses. The DREAM sequence estimates the FA from a single-shot stimulated echo acquisition mode (STEAM) image [28] normalized by an image of the free induction decay (FID), both which are read out by one consecutive RF pulse train. This sequence is fast, has low SAR, but has a relatively narrow dynamic range of 20–50° [30]. In [30], the authors proposed to increase the unbiased B_1_^+^range by acquiring separate B_1_^+^maps with varying nominal preparation angles, thresholding the corresponding FA maps to exclude biased pixels, and then averaging the resulting B_1_^+^maps. That approach was employed in this study by acquiring B_1_^+^maps with three different reference voltages 2/3*V_opt_*, *V_opt_*, and 1.5*V_opt_*). However, it should be noted that for some subject/site combinations (MNI/Subject 1, MSSM/Subject 1, MSSM/Subject 3, NTNU/Subject 1, NTNU/Subject 3) we were unable to acquire the 1.5*V_opt_* scans due to hardware limitations, potentially limiting the dynamic range of the resulting DREAM B_1_^+^maps. In contrast to the DREAM sequence, presaturated turboFLASH (TFL), which measures a series of rapid fast low-angle shot (FLASH) images with different magnitude preparation pulses, has a larger dynamic range but tends to underestimate FA, due to T1 relaxation effects [29]. While it is beyond the scope of this work to investigate the accuracy of DREAM and TFL sequences, one should note that we did observe some differences between the two B_1_^+^mapping methods.

Another point to highlight is that the MPI/NTNU coil pair (MRI.TOOLS) did not always lead to consistent results; B_1_^+^homogeneity and SNR_90_ along the SC showed some differences between the two sites. Some explanations for this may be the fact that (1) different landmarks were used for the *V_opt_* calibration, (2) at both sites the calibrated *V_opt_* exceeded the hardware limit but each site set the final *V_opt_* to something different, (3) it is possible that this coil is more sensitive to slight differences in positioning (along the H/F and/or A/P directions) than other coils.

There is no “one size fits all” when it comes to purchasing or designing and building an MRI coil. We found that the Rapid Biomedical coil (CRMBM/UCL) had the highest B_1_^+^efficiency. The NeuroPoly coil (MGH/MNI) had the highest SNR and greatest coverage along the SC and potential for acceleration (along the F/H and A/P directions). The MRI.TOOLS coil (MPI/NTNU) on the other hand had the most uniform B_1_^+^profile along the cervical SC and acceleration potential along the L/R direction, however, it was limited in its ability to provide the requested flip angles (especially for larger individuals). The latter was also the case for the MSSM coil. Readers interested in the coils’ acceleration capabilities along other combinations of axes can find this data on the project’s open data repositories. No limitations were encountered due to SAR levels for any of the sequences included in our acquisition protocol, for any of the coils.

Given the strengths and weaknesses of each coil, users are advised to select a coil that best suits their unique needs. For example, users seeking to acquire power-intensive sequences may be swayed by the high B_1_^+^efficiency of the Rapid Biomedical coil, likely conferred by its large tightly fitting coil elements. Others interested in acquiring images that provide good coverage along the A/P, L/R, and H/F directions may prefer the NeuroPoly coil design with its large transmit region and its broadly distributed receive elements. The NeuroPoly coil could be useful for acquiring high-resolution anatomical and functional scans considering large FOVs. The acceleration potential of different coils should also be considered, as it enables faster image acquisitions and reduced susceptibility distortions when using EPI sequences (often used for diffusion and fMRI applications). The MRI.TOOLS coil, with many small receive elements distributed around the neck, is attractive for high-resolution imaging of the cervical SC with high acceleration capabilities along L/R (and/or A/P) directions. If greater coverage along the length of the SC is desired, the NeuroPoly coil has a greater promise for acceleration.

## Acknowledgments

We thank Marcus Couch who provided useful suggestions, Mathieu Boudreau who helped host the Jupyter notebooks,Helmar Waiczies for his description of the MRI.TOOLS coil, and Philipp Ehses who provided the 2D DREAM sequence. This research was undertaken thanks, in part, to funding from NTNU Biotechnology, the Canada First Research Excellence Fund (TransMedTech Institute), the Natural Sciences and Engineering Research Council of Canada, the Canada Research Chair in Quantitative Magnetic Resonance Imaging, the Canadian Institute of Health Research, the Canada Foundation for Innovation, the Fonds de Recherche du Québec - Santé, Polytechnique Montreal, and the Quebec Bioimaging Network (QBIN), and MITACS Accelerate Fellowship. Imaging performed at the Athinoula A. Martinos Center for Biomedical Imaging used resources provided by the Center for Functional Neuroimaging Technologies (P41EB015896) and the Center for Mesoscale Mapping (P41EB030006), Biotechnology Resource Grants supported by the National Institute of Biomedical Imaging and Bioengineering, National Institutes of Health (NIH). The NIH also provided support through grants S10OD023637, R01EB027779 and R21EB031211. This research was also supported in part by the MGH/HST Athinoula A. Martinos Center for Biomedical Imaging, as well as by the Centre National de la Recherche Scientifique (CNRS). The content is solely the responsibility of the authors and does not necessarily represent the official views of the NIH. Data acquisition at the Max Planck Institute for Human Cognitive and Brain Sciences was supported by the Max Planck Society and the European Research Council (under the European Union’s Horizon 2020 research and innovation programme; grant agreement No 758974).

## Data Availability Statement

All data analysis code and protocol PDF printouts and executable (.exar1) files, and SOPs can be found on our projects GitHub organization: https://github.com/spinal-cord-7t

Phantom and human data are located here: https://openneuro.org/datasets/ds005090 and here: https://openneuro.org/datasets/ds005025.

## Disclaimers

Since January 2024, Dr. Barry has been employed by the National Institute of Biomedical Imaging and Bioengineering at the NIH. This work was co-authored by Robert Barry in his personal capacity. The opinions expressed in this study are his own and do not necessarily reflect the views of the NIH, the Department of Health and Human Services, or the United States government. Since September 2023, Daniel Papp has been an employee of Siemens Healthineers Sweden AB, but was a postdoctoral researcher at NeuroPoly Lab during the time of the study and initial preparation of this abstract. This work was co-authored by Daniel Papp in his personal capacity, and no company resources were used. Opinions expressed in this study do not necessarily reflect the view of Siemens Healthineers Sweden, or Siemens Healthcare.

