## Supplementary Materials for "Multi-center benchmarking of cervical spinal cord RF coils for 7 T MRI: A traveling spines study"

| Sequence | Resolution ($mm^3$) | FOV ($mm^2$) | Slices | TR (ms) | TE (ms) | FA | Voltage | TA |
|---|---|---|---|---|---|---|---|---|
| **Sagittal TFL** | 2.4x2.4x2.5 | 340x208 | - 2 slice groups of 28 slices<br>- 2.5 mm slice thickness and 2.5 mm slice gap | 34420 | 2.31 | - pre-sat FA= 90° (80° for Magnetom sites)<br>- excitation FA= 10° | reference voltage calibrated by the scanner | 1m10s |
| **Sagittal TFL** | 2.4x2.4x2.5 | \|\| | \|\| | \|\| | \|\| | \|\| | $V_{opt}$ | 1m10s |
| **Sagittal DREAM** | 4.1x4.1x2.5 | 384x178 | 11 slices of 2.5 mm slice thickness | 6000 | TE1/TE2 = 2.04/3.18 | FA1/FA2 = 50/6° | \|\| | 1m08s |
| **Sagittal DREAM** | 2.5x2.5x2.5 | 200x200 | \|\| | \|\| | \|\| | \|\| | \|\| | 1m08s |
| **Sagittal DREAM** | 2.5x2.5x2.5 | \|\| | \|\| | \|\| | \|\| | \|\| | $(2/3) * V_{opt}$ | 1m08s |
| **Sagittal DREAM** | 2.5x2.5x2.5 | \|\| | \|\| | \|\| | \|\| | \|\| | $1.5 * V_{opt}$ or $V_{HW,limit}$ (whichever was lower) | 1m08s |

**Supplementary Table 1:** Acquired $B_1^+$ mapping sequences and their parameters. "||" indicates that the field is identical to that of the above-row, $V_{opt}$ is the optimal reference voltage, and $V_{HW,limit}$ is the hardware limit voltage for a given coil.

| Sequence | Resolution ($mm^3$) | FOV ($mm^2$) | Slices | TR (ms) | TE (ms) | FA | Phase Encode | Voltage | TA |
|---|---|---|---|---|---|---|---|---|---|
| **Sagittal coilQA** | 1.5x1.5x2.0 | 384x384 | 13 slices of 2.0 mm slice thickness with 20% slice gap | 30 | 6 | 12° | H>F | $V_{opt}$ | 3m20s |
| **Sagittal coilQA** | 1.5x1.5x2.0 | 192x192 | \|\| | \|\| | \|\| | \|\| | \|\| | \|\| | 1m40s |
| **Axial coilQA** | 0.5x0.5x5.0 | \|\| | 7 slices of 5.0 mm slice thickness with 300% slice gap | \|\| | \|\| | \|\| | R>L | \|\| | 2m41s |
| **Sagittal 3D GRE** | 2.0x2.0x2.0 | 384x288 | 44 slices of 2.0 mm slice thickness without slice gap | 550 | 3.8 | 30° | R>L | \|\| | 1m21s |

**Supplementary Table 2:** Acquired $B_1^-$ mapping sequences and their parameters. "||" indicates that the field is identical to that of the above-row, and $V_{opt}$ is the optimal reference voltage.

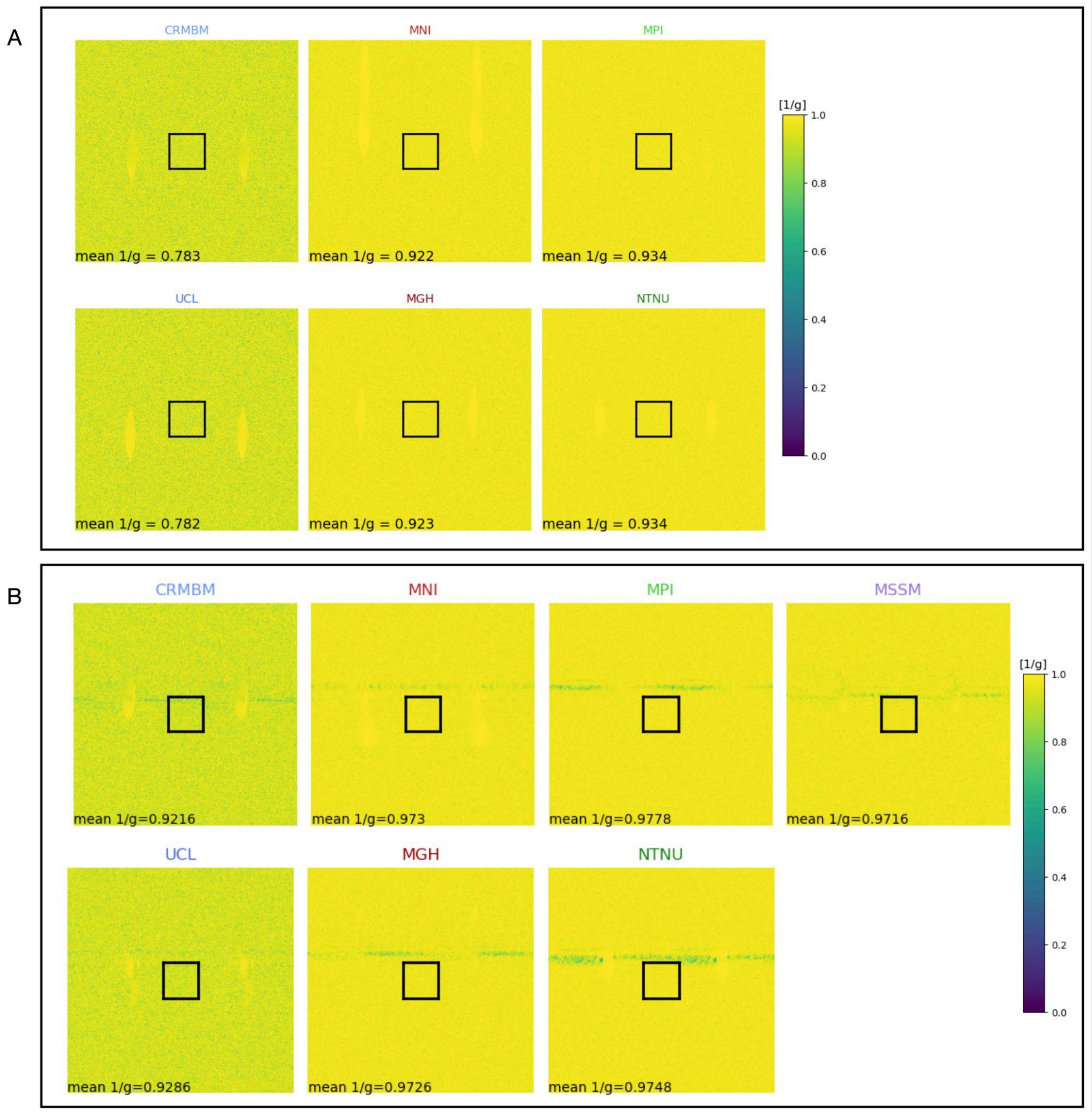


**Supplementary Figure 1:** (A) 1/g maps with $R_{L/R}$ = 2 of Spinoza at each site, except for MSSM. (B) 1/g factor maps with $R_{L/R}$ = 2 for a single representative subject across all sites (Subject 2).

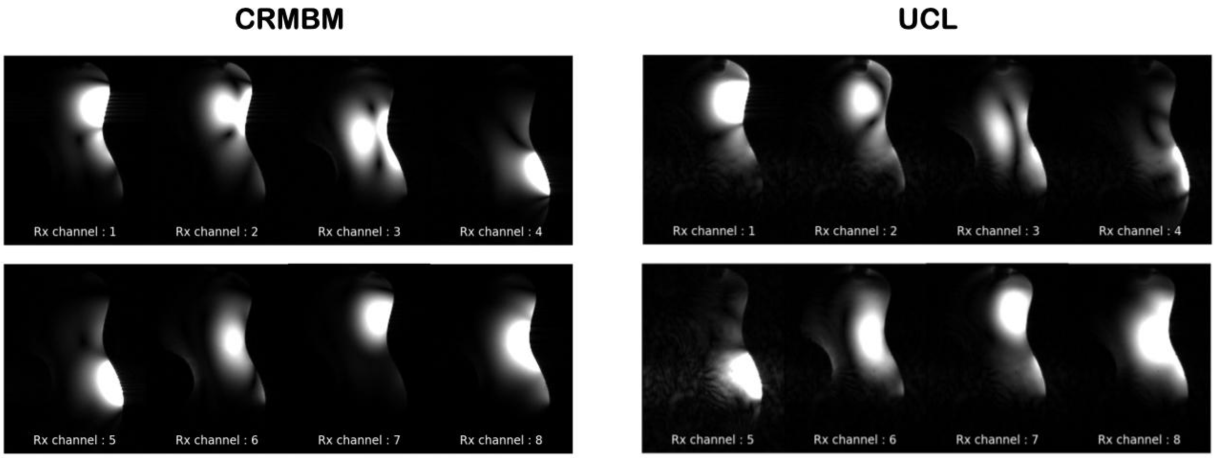


**Supplementary Figure 2:** Individual channel GRE phantom sagittal images for CRMBM and UCL (Subject 2).

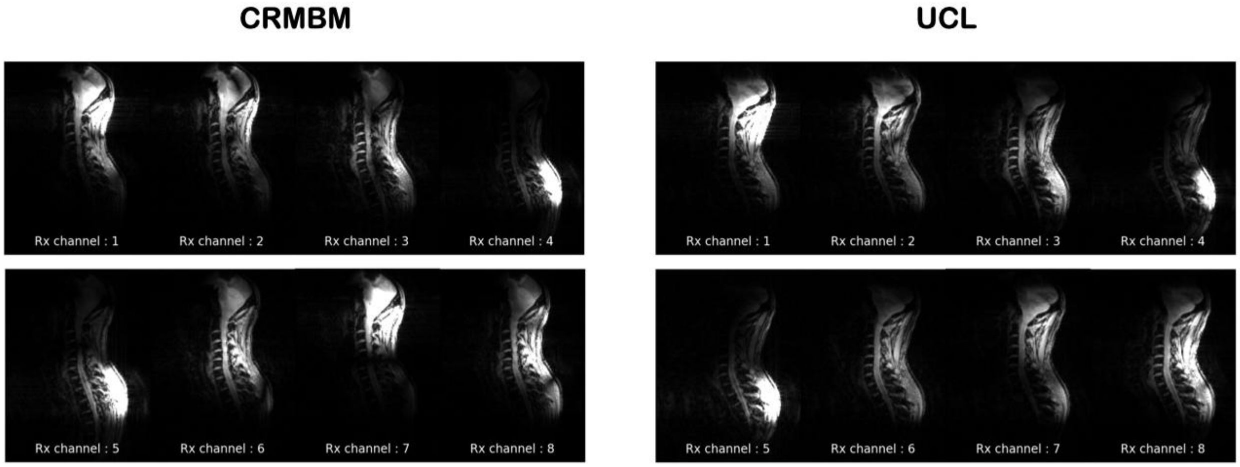


**Supplementary Figure 3:** Individual channel GRE in-vivo sagittal images for CRMBM and UCL (Subject 2).

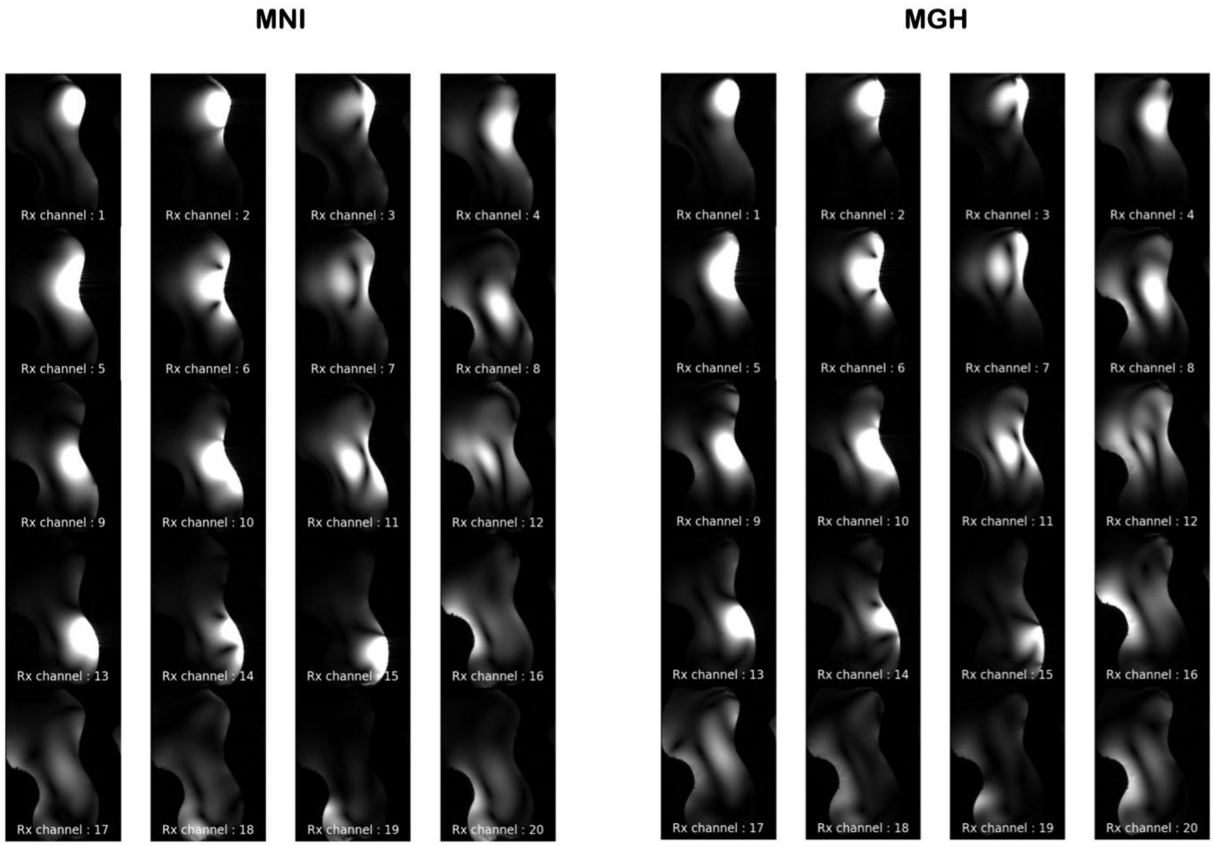


**Supplementary Figure 4:** Individual channel GRE phantom sagittal images for MNI and MGH (Subject 2).

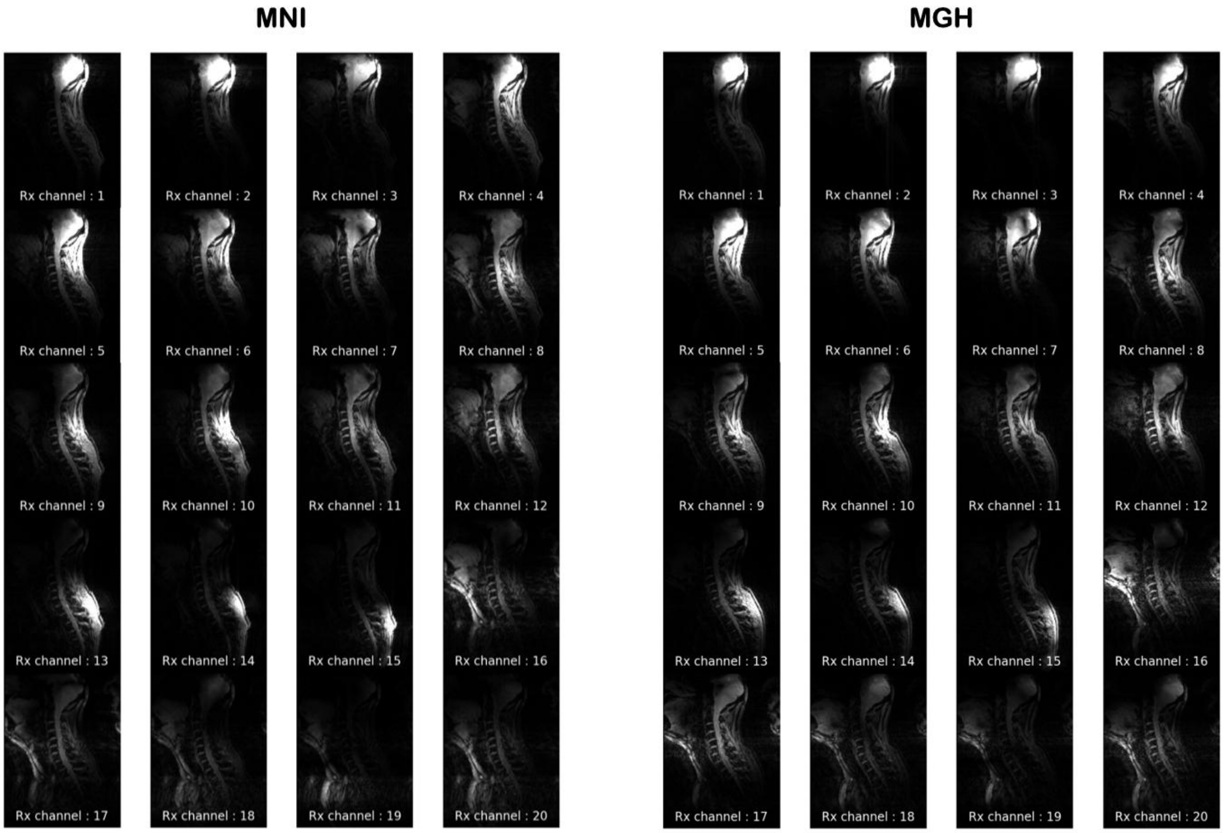


**Supplementary Figure 5:** Individual channel GRE in-vivo sagittal images for MNI and MGH (Subject 2).

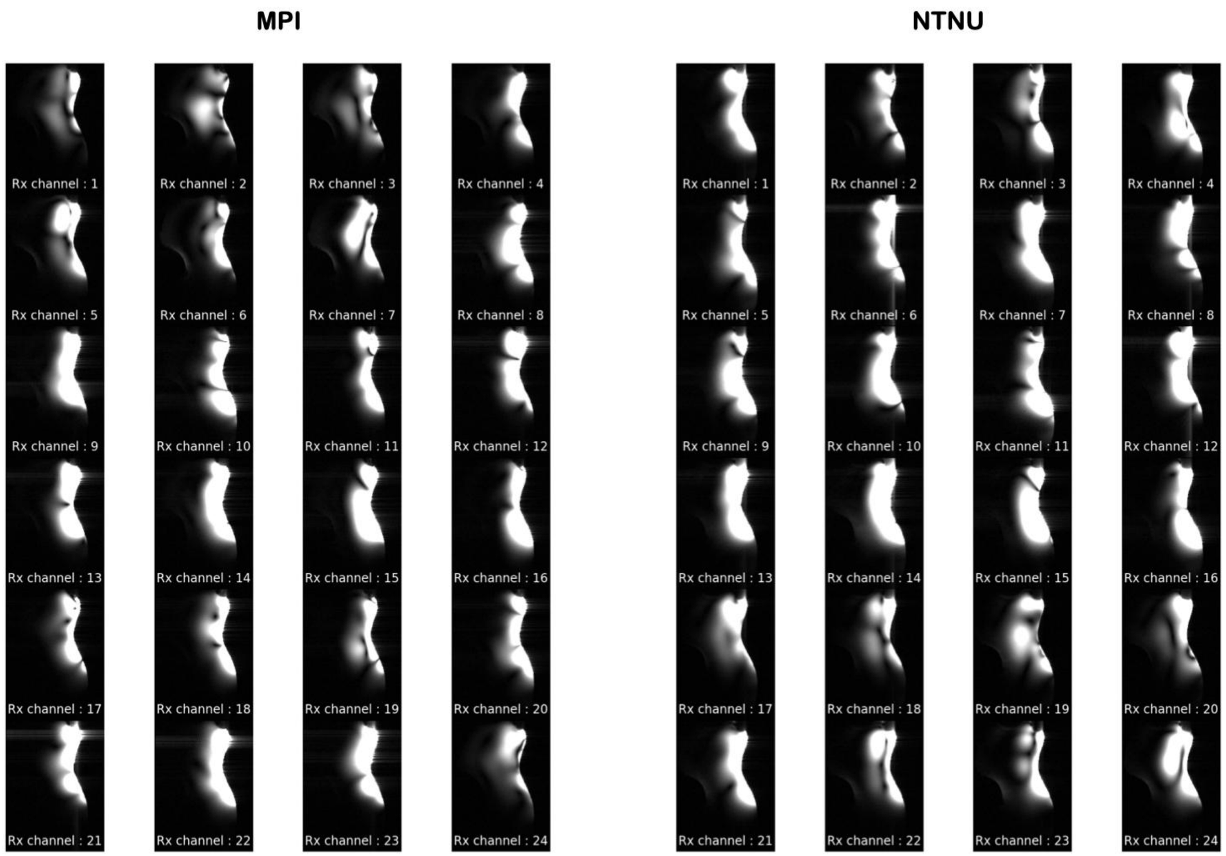


**Supplementary Figure 6:** Individual channel GRE phantom sagittal images for MPI and NTNU (Subject 2).

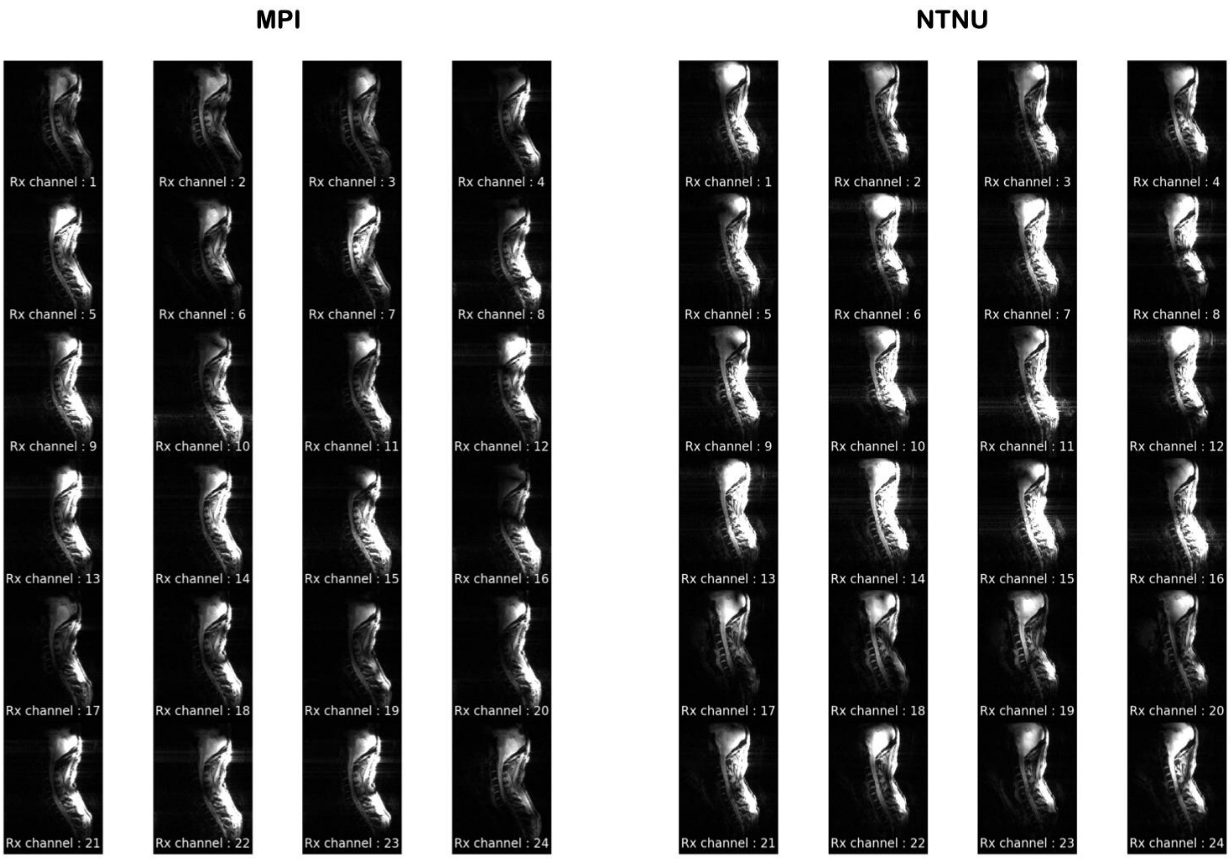


**Supplementary Figure 7:** Individual channel GRE in-vivo sagittal images for MPI and NTNU (Subject 2).

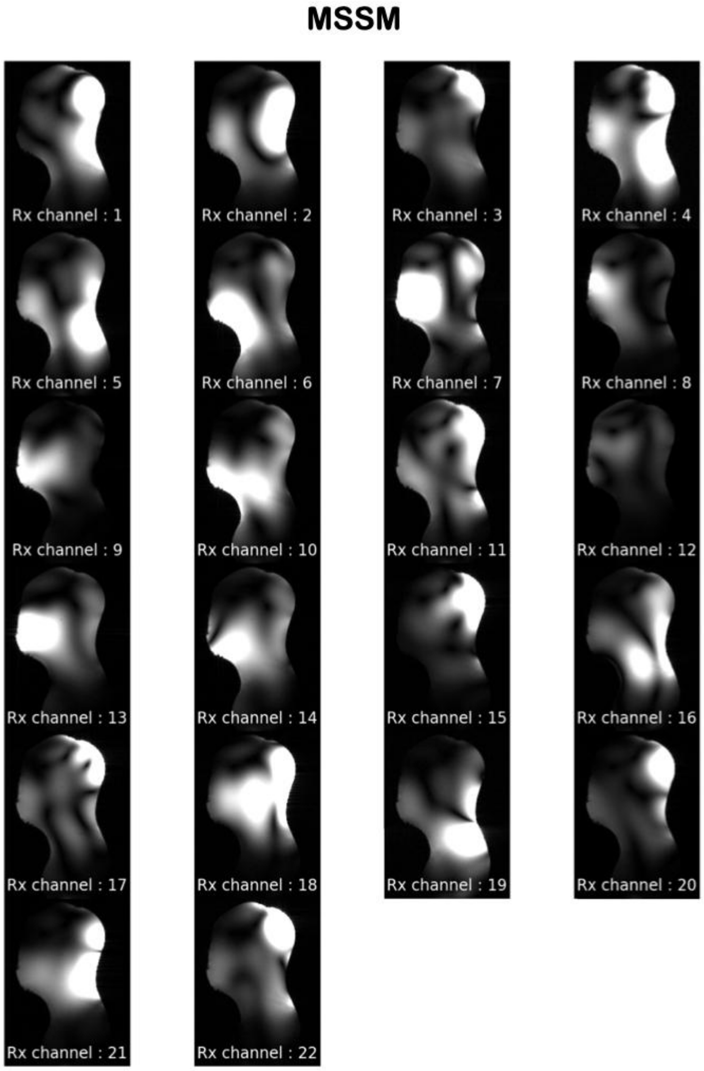


**Supplementary Figure 8:** Individual channel GRE phantom sagittal images for MSSM (Subject 2).
